# Kynurenine Metabolism Mediates Tumor Progression in Renal Cell Carcinoma

**DOI:** 10.64898/2026.02.10.705186

**Authors:** Kayla M. Miller, Skye Rounseville, Raul Castro-Portuguez, Robert Railey, Hope Dang, Kailyn Dundore, Luis Espejo, Samuel Freitas, George L. Sutphin

## Abstract

Renal cell carcinoma (RCC) is characterized by dysregulation of the kynurenine pathway (KP), which converts tryptophan to NAD^+^ while generating immunomodulatory metabolites. Therapeutic efforts have focused on inhibiting IDO1 at the pathway entry point, but the functional consequences of targeting downstream KP enzymes remain poorly characterized. We used CRISPR/Cas9 to generate knockouts of three KP enzymes in the RENCA murine RCC model—kynurenine 3-monooxygenase (KMO), quinolinic acid phosphoribosyltransferase (QPRT), and 3-hydroxyanthranilic acid dioxygenase (HAAO)—and evaluated effects on cell migration, colony formation, tumor burden, metastasis, and survival. KMO and QPRT knockouts consistently reduced migratory capacity and colony size *in vitro*. However, *in vivo* effects were distinct: while QPRT knockout reduced tumor burden, KMO knockout did not. Notably, we did not detect metastasis in female mice receiving KMO knockout RENCA cells. HAAO knockout produced divergent effects, increasing migration and colony size *in vitro*, but reducing tumor burden and metastasis *in vivo*. Mice challenged orthotopically with all three knockout cell lines had significantly extended survival compared to mice receiving wild type cells. These results indicate that individual KP enzymes exert distinct, context-dependent effects on RCC progression. The enhanced *in vitro* aggressiveness coupled with reduced *in vivo* tumorigenicity observed in HAAO knockout RENCA cells illustrates that cell culture phenotypes do not reliably predict tumor behavior, particularly when perturbing metabolic pathways with pleiotropic effects. Our findings suggest that targeting specific KP enzymes warrants further investigation as a therapeutic strategy in RCC.

## Introduction

Homeostatic cellular metabolism requires a balance between energy production and macromolecule synthesis to support cell growth and function. In cancer cells, this equilibrium is disrupted, and altered cellular metabolism is a recognized hallmark of cancer that helps sustain rapid tumor cell proliferation^1,2^. The metabolic alterations in Renal Cell Carcinoma (RCC) are particularly interesting, as this subtype of tumors exhibit distinct metabolic profiles compared to normal kidney tissue^3^. RCC is characterized by a reliance on aerobic glycolysis, similar to the Warburg effect observed in other cancer types ^4,5^.

The kynurenine pathway (KP; Fig. 1A) is increasingly recognized as a metabolic vulnerability in cancer, and specifically in RCC where KP dysregulation is frequently observed^6^. The most common framing is that tryptophan catabolism through the KP confers two major advantages to tumor cells. First, depletion of tryptophan in the tumor microenvironment suppresses antitumor immunity. Tryptophan shortage can activate the stress kinase GCN2 in T cells, leading to cell cycle arrest and anergy^7^, and suppress mTOR signaling, impairing effector T cell function^8^, though the *in vivo* relevance of GCN2-mediated suppression has been challenged by data from some tumor models^9,10^. Kynurenine and downstream metabolites can also activate the aryl hydrocarbon receptor (AhR)^11^, promoting regulatory T cell differentiation and further dampening immune responses^12–14^. Second, the KP is also the de novo route for synthesizing nicotinamide adenine dinucleotide (NAD^+^), an essential cofactor for glycolysis, the tricarboxylic acid cycle, and oxidative phosphorylation. NAD^+^ also serves as a substrate for poly(ADP-ribose) polymerases (PARPs) and sirtuins, which mediate DNA repair, chromatin remodeling, and metabolic regulation—processes critical for cancer cell survival under genotoxic and metabolic stress^15,16^. These mechanisms position the KP as a pathway that promotes both immune evasion and the bioenergetic needs of rapidly proliferating tumor cells.

**Figure 1:**
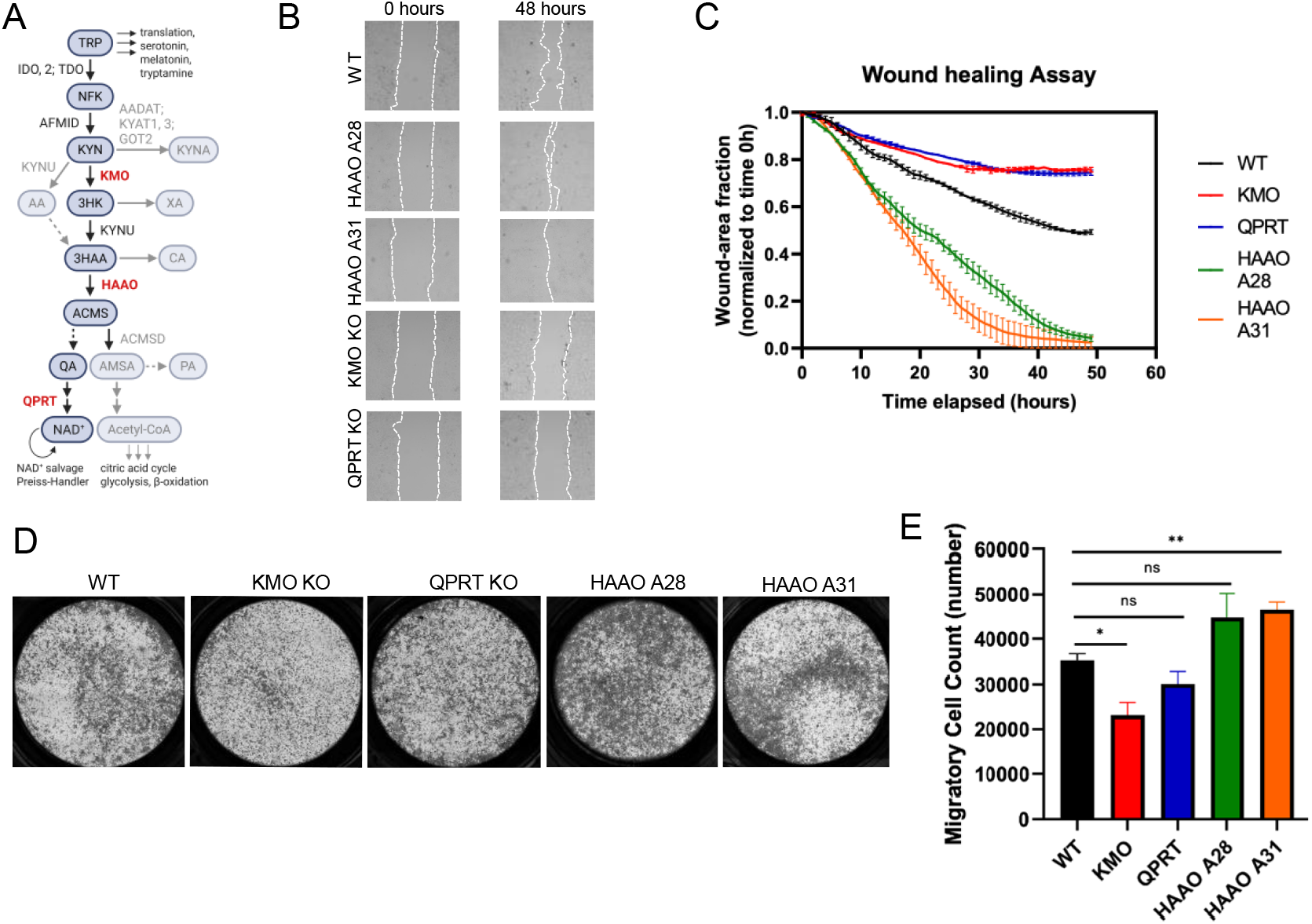
HAAO Knockout Increases Migratory and Invasive Potential in RENCA Cells. (A) Schematic or kynurenine metabolic pathway. (B) Representative images of transformed RENCA cells at 0h and 48h following scratch wound assay. (C) Quantification of wound healing time. (D) Representative images of transwell invasion assay of RENCA cell lines at 48 hours. (E) Quantification of transwell assay migratory cell count (D) at 48 hours. Line chart represents mean at the indicated time point. Bars represent mean. Error bars represent SEM. * p < 0.05, ** p < 0.001, by Welch’s t test with the Holm-Šídák multiple test correction.

However, this framework treats the KP as a functionally uniform pathway. The KP has multiple branch points where different enzymes control the accumulation of metabolites with distinct biological properties^17^. While most therapeutic efforts have focused on inhibiting IDO1 to restore tryptophan availability, the functional consequences of targeting downstream KP enzymes remain incompletely characterized^17,18^. In this study, we focus on three enzymes: kynurenine 3-monooxygenase (KMO), which initiates the hydroxylation branch leading to 3-hydroxykynurenine production; quinolinic acid phosphoribosyltransferase (QPRT), which catalyzes the final step in de novo NAD+ biosynthesis; and 3-hydroxyanthranilic acid dioxygenase (HAAO), which converts 3-hydroxyanthranilic acid (3HAA) to quinolinic acid (QA, Fig. 1A). We predicted that genetic ablation of these enzymes would produce different metabolic consequences. All three knockouts are predicted to impair de novo NAD+ synthesis, but result in accumulation of distinct intermediate metabolite species: kynurenine (KYN) in KMO knockouts, an aryl hydrocarbon receptor ligand with immunosuppressive properties; 3HAA in HAAO knockouts, a redox-active metabolite with immunomodulatory properties^19–21^; and QA in QPRT knockouts, an N-methyl-D-aspartic acid receptor (NMDA) receptor agonist that can induce cellular stress^22,23^. Previously, we found that genetic reduction of HAAO or supplementation with exogenous 3HAA extends lifespan in both *C. elegans* and mice^24^.

Whether blocking individual KP enzymes produces distinct effects on tumor progression remains unknown.

As a redox-active compound, 3HAA can function as both an antioxidant and a pro-oxidant depending on local concentration and cellular redox status^19^. In immune contexts, 3HAA suppresses T cell proliferation through multiple mechanisms, including glutathione depletion in activated T cells and inhibition of dendritic cell activation^20,25^. However, 3HAA also exhibits anti-inflammatory properties under certain conditions, inducing expression of hemeoxygenase-1 and other cytoprotective enzymes in astrocytes and promoting a shift from pro-inflammatory to regulatory immune phenotypes^21^. Because 3HAA can both induce oxidative stress and activate antioxidant responses, 3HAA accumulation following HAAO inhibition could produce different outcomes than interventions targeting earlier or later pathway steps. In cancer, where both immune suppression and oxidative stress resistance contribute to tumor progression, the net effect of elevated 3HAA may depend on the tumor microenvironment and the metabolic state of both tumor cells and infiltrating immune cells^17,18,26,27^.

To investigate whether individual KP enzymes have distinct roles in RCC progression, we generated knockout cell lines for *Kmo, Qprt*, and *Haao* using CRISPR/Cas9 gene editing in the RENCA murine RCC model. We hypothesized that disruption of these enzymes would produce enzyme-specific effects on tumor cell behavior and tumor progression, reflecting the distinct metabolic and signaling functions of the metabolites they control. Using a combination of *in vitro* cellular assays to assess migratory capacity and clonogenic growth, and *in vivo* syngeneic tumor models to evaluate tumor burden, metastasis, and host survival, we characterized the functional consequences of blocking each enzyme. We found that knockout of *Haao* produces effects distinct from *Kmo* or *Qprt*, with increased aggressiveness *in vitro* but reduced tumorigenicity *in vivo*, while all three knockouts improved survival. These results demonstrate that the KP comprises distinct metabolic nodes with enzyme-specific and context-dependent effects on cancer progression.

## Materials and Methods

### Cell Lines

Murine renal carcinoma cell lines were maintained in RPMI (GIBCO) supplemented with 10% fetal bovine serum (FBS, Thermo Fisher Scientific and Gemini Bio Products),1% penicillin streptomycin (PenStrep, Gibco) 1% NEAA, 1% Glutamax, 1% Sodium Pyruvate, and Puromycin (1 ng/uL) at 37°C under 5% CO_2_. Experiments were conducted on cells maintained between passage 3 and passage 10 to ensure stability and reduce the risk of phenotypic drift. All cell lines were tested for mycoplasma and were authenticated by the Experimental Mouse Shared Resource (EMSR) at The University of Arizona.

### Colony Formation (Clonogenic Assay)

Cells were first trypsinized and counted. 200 cells per well were seeded in 6 well plates and cultured for 14 days. Formed cellular colonies were fixed with 4% paraformaldehyde (PFA) for 15 minutes and the stained with 1% crystal violet. Colonies containing greater than or equal to 50 cells were counted.

### Migration

Cell migration was assessed using previously established methods^28^. Cells were seeded at 1×10^5^ cells on transwell plates (8 μm pore PET membrane, Corning Cat#3464). The top chamber contained serum free media; the bottom chamber contained 10% FBS. After 48 hours given for migration, the top of the chamber was scraped with a cotton swab, and inserts were fixed for 10 minutes in 4% PFA. Cells were stained for 10 minutes with 1% crystal violet, and the number of migrated cells counted using FIJI image analysis.

### Wound Healing

Cells were seeded and grown to confluency in 4 Chamber Glass Bottom dishes (CellVis). A horizontal scratch was made using a 10 μL pipette tip^29^. Cells were imaged every hour for 48 hours using an Olympus IX83 confocal microscope equipped with an environmental chamber controlling temperature, atmosphere (5% CO2), and humidity (95%). Images were analyzed using the custom entropy_DIC script developed in our lab using MATLAB (https://github.com/Sam-Freitas/entropy_DIC).

### Generation of RENCA Cell Lines

Transgenic cell lines were generated by the University of Arizona Genome Editing Facility. *Kmo* exons 2 and 3, *Qprt* exon 2, HAAO exon 1 were deleted using CRISPR/Cas9 to make double-strand breaks on either side of exon. Genotype of each mutant cell clone was verified using PCR.

### RENCA Renal Cancer Mouse Model

Sixty-four eight-week-old BALB/cByJ mice (32 male, 32 female) were acquired from The Jackson Laboratory and allowed a two-week acclamation period. The survival study involved orthotopically injecting control (WT), KMO-/-, HAAO-/-, and QPRT-/- RencaLuc cells into the left kidneys of each mouse on Day 0. Tumor growth was monitored using *in vivo* bioluminescence at days 3, 10, 17, 24, and 31. Mice were observed at least twice a week during the first two weeks and daily from Day 14 onward. Mice meeting the criteria for moribundity were euthanized, and gross necropsy was performed to evaluate primary and metastatic tumor burden, as well as any other pathology. All surviving mice were euthanized on Day 60 for gross necropsy.

### Bioluminescence

To evaluate live *in vivo* tumor progression, D-luciferin (150 mg/kg body weight) was administered via intraperitoneal injections and mice were subjected to Bioluminescence Imaging (BLI) with a Spectral Instruments Imaging Lago at the University of Arizona Translational Bioimaging Resource.

### Data Analysis

All graphs were generated using GraphPad Prism software (version 8. 4.3.686) unless otherwise stated. Survival between groups of mice was compared by a log rank test with Holm-Šídák multiple test correction. No animals were censored from survival studies with the exception of the mice still alive at the end of the study period. Comparisons between groups in all other experiments were calculated by unpaired t test (two-sided) with Welch’s correction and a Holm-Šídák multiple test correction, with alpha = 0.05. Significance is annotated with asterisks as follows: * p < 0.05, ** p < 0.01, *** p < 0.001, and **** p < 0.0001. Error bars represent the standard error of the mean (SEM).

## Results

Dysregulation of the kynurenine pathway is a ubiquitous feature of human RCC and has been implicated in tumor immune evasion and metabolic adaptation, making this pathway an attractive target for therapeutic intervention^6,13,17^. To test how disrupting individual enzymes in the NAD^+^-producing branch of the kynurenine pathway influences RCC progression, we used the murine RENCA cell line, a well-established RCC model derived from a spontaneous tumor in BALB/c mice that supports orthotopic implantation into the kidney of immunocompetent hosts and recapitulates primary tumor growth and spontaneous metastasis^30–32^. Using CRISPR/Cas9, we generated clonal RENCA lines with targeted knockout of *Kmo, Haao*, or *Qprt* and compared their behavior *in vitro* and in an orthotopic syngeneic tumor model *in vivo*.

Cell migration is a critical process in cancer metastasis and is influenced by metabolic state and cytoskeletal dynamics. Prior studies have shown that metabolites such as KYN and 3HAA can affect redox signaling and modulate migration-related gene expression^18,33,34^.To test how specific KP enzymes influence these behaviors, we conducted wound healing and transwell assays in our *Kmo, Qprt*, and *Haao* knockout RENCA cell lines. Wound healing assays showed that *Haao* knockout cells closed the artificial wound more rapidly than wild type (WT) cells (Fig. 1B, C), which may reflect increased migration, proliferation, or both. Conversely, *Kmo* and *Qprt* knockouts showed delayed wound healing, suggestive of impaired motility and/or proliferation. To specifically evaluate migratory potential, we next conducted a transwell assay. *Haao* knockout cells migrated more readily through the membrane pores, while *Kmo* and *Qprt* knockouts displayed reduced migration (Fig. 1D, E). These results suggest that HAAO loss confers a pro-migratory and pro-invasive phenotype, potentially linked to the accumulation of 3HAA, which has known redox signaling properties.

Proliferation and clonogenic survival are essential hallmarks of tumorigenic potential. Cancer cells with enhanced colony formation often possess greater resistance to environmental stress and increased proliferative capacity^35^. To determine if the observed differences in wound closure was driven by changes in cell division rates, we performed a proliferation assay. Direct cell counting over a 96-hour period showed that *Haao* and *Qprt* knockout cells proliferated at a rate comparable to WT controls (Fig. 2A), while *Kmo* knockouts trended toward reduced proliferation without reaching statistical significance. Despite having proliferation rates similar to WT, *Haao* knockout cells formed colonies with significantly larger surface areas in clonogenic assays (Fig. 2B, C). Since cell division rates are unchanged in *Haao* knockouts, this increased colony footprint likely reflects the enhanced migratory and spreading phenotype observed in our wound healing and transwell assays. Conversely, *Kmo* and *Qprt* knockouts formed smaller colonies, consistent with their reduced migratory potential and, in the case of KMO, potentially also a reduced proliferation. The number of colonies formed was similar across genotypes (Fig. 2C, D).

**Figure 2:**
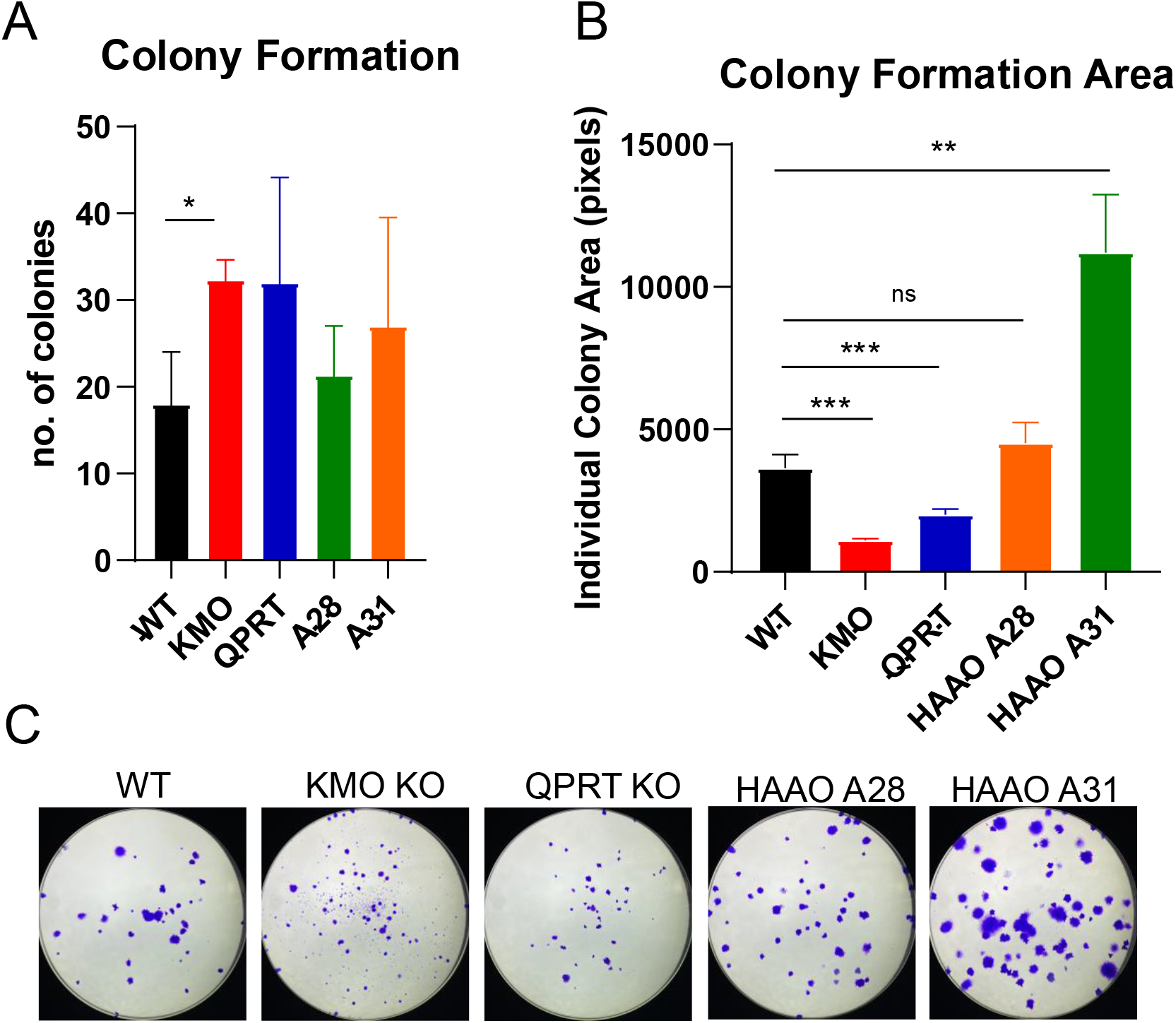
Colony Formation Assay Reveals Altered Proliferative Behavior. (A) Total number of colonies formed and (B) colony area for each RENCA KP knockout cell line. (C) Representative images of colonies for each cell line visualized by crystal violet staining. Bars indicate mean and error bars SEM. * p < 0.05, ** p < 0.001, *** p < 0.0001, by Welch’s t test with the Holm-Šídák multiple test correction.

The *in vivo* tumor microenvironment introduces additional complexity, including immune surveillance, stromal interactions, and nutrient availability. Previous work indicates that tryptophan catabolism contributes to immune tolerance and NAD^+^ availability in the tumor niche^17^. We evaluated the impact of *Kmo, Qprt*, or *Haao* knockout on tumor growth and metastasis using a RENCA syngeneic mouse model. Unexpectedly, *in vivo* tumor burden (Fig. 3A) was lowest in the *Haao* knockout group despite their high migratory and proliferative behavior *in vitro*. This contradiction may reflect effects of HAAO loss that are not captured *in vitro*—such as altered redox balance or immune interactions, potentially mediated by 3HAA accumulation—which could suppress tumor establishment and expansion *in vivo*. Bioluminescence imaging revealed fewer and less severe metastatic events in *Haao* knockout mice (Fig. 3B), supporting the idea that *in vitro* motility does not guarantee metastatic success, especially in the absence of immune evasion and stromal adaptation. Notably, we did not observe any metastatic events in female mice receiving *Kmo* knockout RENCA cells.

**Figure 3:**
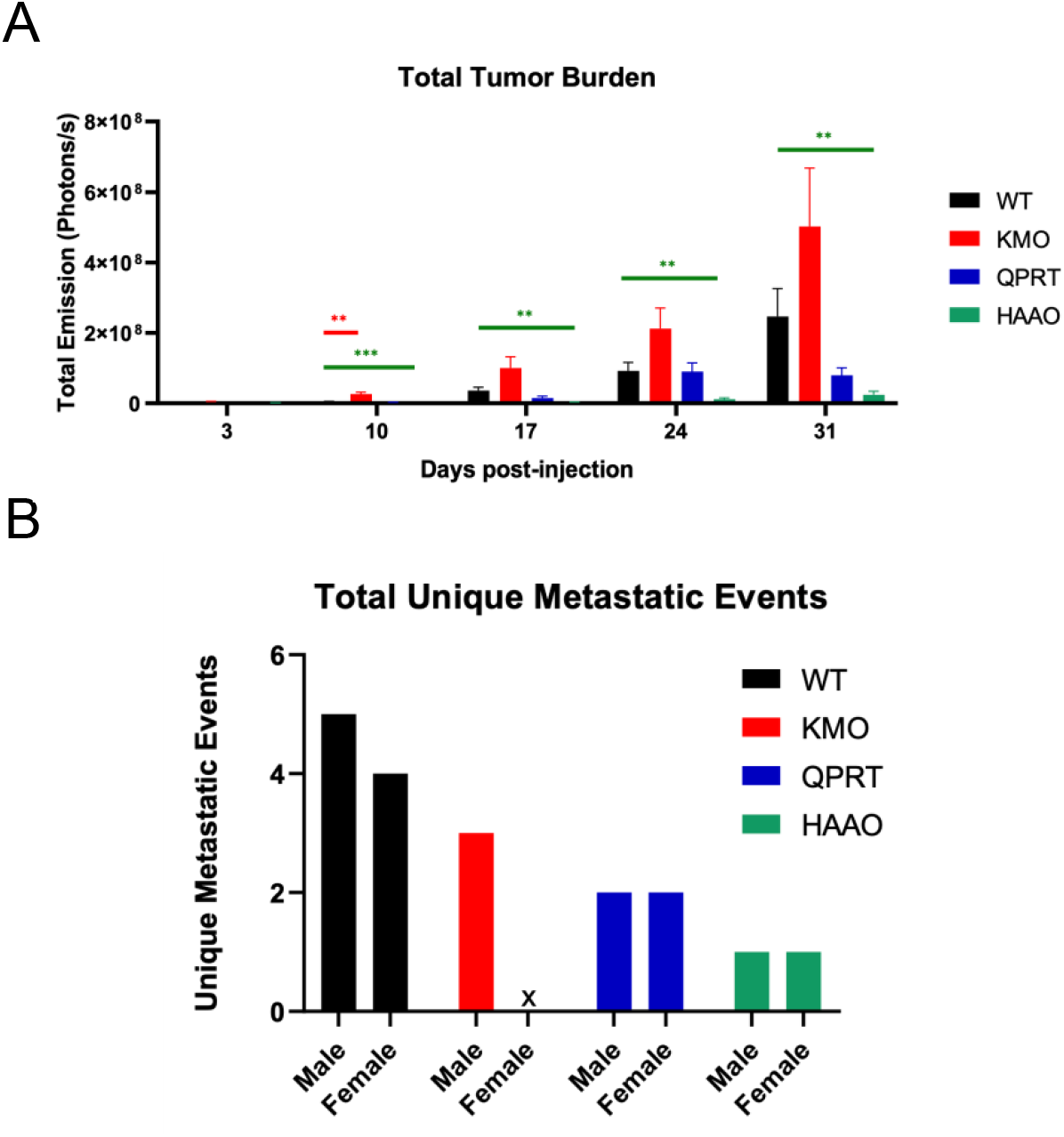
HAAO Knockout Reduces Tumor Burden and Metastasis. (A) Total tumor burden per condition quantified by bioluminescent imaging. All statistically significant differences indicated by stars. (B) Total unique metastatic events observed via bioluminescence imaging throughout the duration of the study. “x” indicates 0 observed metastatic events. * p < 0.05, ** p < 0.01, *** p < 0.001, by Welch’s t test with the Holm-Šídák multiple test correction.

Survival outcomes in cancer are influenced by both tumor burden and the systemic effects of metabolic remodeling. NAD^+^ depletion has been shown, for example, to impair DNA repair and cellular homeostasis, leading to increased sensitivity to stress and reduced tumor viability^15^. We next determined whether disrupting the KP could improve survival outcomes *in vivo*. Mice receiving orthotopic injection of all three KP knockout RENCA cell lines had improved survival relative to WT (Fig. 4). These results suggest that disrupting enzymes in the oxidative branch of the KP can slow tumor progression and improve outcomes *in vivo*. The consistent trend across all knockouts suggests that targeting several KP enzymes may have therapeutic potential.

**Figure 4:**
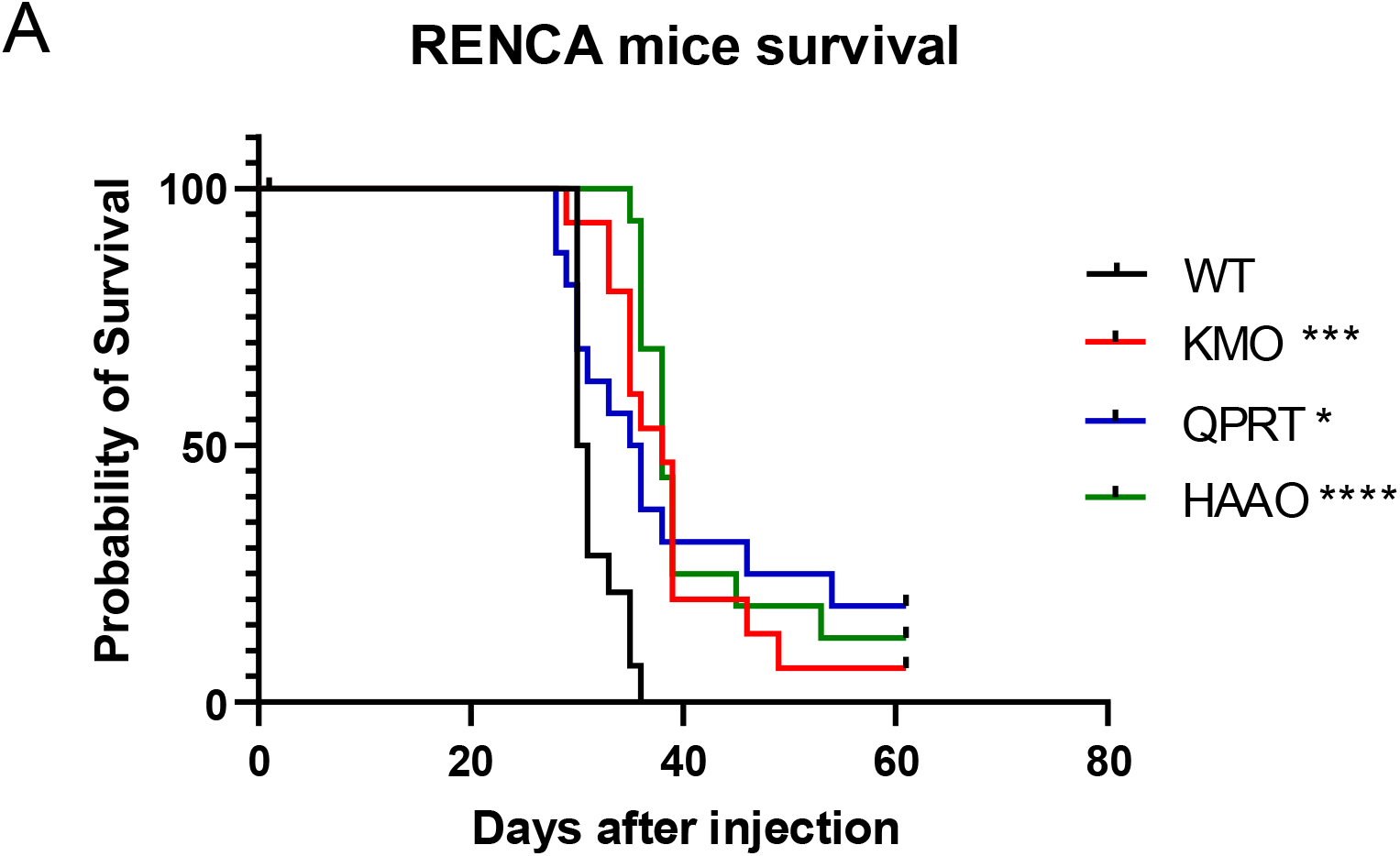
Knockouts of KP Enzymes Improve Overall Survival. Kaplan-Meier survival curves for male mice injected orthotopically with indicated RENCA KP knockout cell lines (n = 16 mice per condition). * P < 0.05, *** P < 0.0001, **** P < 0.0001 by the log rank test with the Holm-Šídák multiple test correction

## Discussion

The development of cancer therapies targeting the KP has focused largely on inhibiting IDO1 to restore tryptophan availability and reduce immunosuppression. More recent attention has expanded to include downstream enzymes and intermediate metabolites, though the functional consequences of targeting specific pathway steps remain incompletely characterized. Our findings demonstrate that genetic disruption of three KP enzymes—KMO, QPRT, and HAAO—produces distinct effects on tumor cell behavior and tumor progression, suggesting that the KP contains metabolic nodes with enzyme-specific functions in RCC. *Haao* knockout increased cell migration and colony area *in vitro* but reduced tumor burden and metastasis *in vivo*. This is consistent with a model in which reduced HAAO activity enhances motility *in vitro* but, *in vivo*, creates vulnerabilities to immune surveillance and/or oxidative stress. However, we did not measure immune infiltration or redox status, so this remains speculative. In contrast, *Kmo* knockouts presented a different uncoupling of phenotypes. *Kmo* knockout cells trended toward reduced proliferation and displayed reduced clonogenicity *in vitro*, yet trended toward an increased primary tumor burden *in vivo. Qprt* knockout cells displayed impaired tumor-promoting phenotypes in both contexts. Mice challenged with all three KP knockout cell lines displayed significantly extended survival relative to mice challenged with unmodified RENCA cells, suggesting that disruption of the NAD^+^-producing branch of the KP can impair RCC progression *in vivo*. These results establish that individual KP enzymes produce distinct effects on tumor behavior and suggest that these differences may reflect the specific metabolites that accumulate at each pathway step and the physiological context in which tumors develop.

The most unexpected finding from this study is the divergent behavior of *Haao* knockout cells *in vitro* versus *in vivo*. In culture, *Haao* knockout cells displayed increased migratory capacity in both wound healing and transwell assays, and formed larger colonies than wild type cells, phenotypes typically associated with more aggressive tumor behavior. However, when implanted into mice, *Haao* knockout cells produced smaller tumors, fewer metastatic events, and longer host survival compared to wild type controls. Such discordances between *in vitro* aggressiveness and *in vivo* tumorigenicity have been reported for other metabolic interventions^36,37^, underscoring that cell culture systems lack immune cells, stromal interactions, and physiological oxygen gradients that critically influence tumor behavior.

HAAO catalyzes the conversion of 3HAA to QA, and based on pathway topology, HAAO knockout is expected to result in 3HAA accumulation. Although we did not directly measure metabolite levels here, the context-dependent biological activities reported for 3HAA could plausibly contribute to the divergent phenotypes. 3HAA can function as both a pro-oxidant and antioxidant depending on local redox conditions^19^, suppress T cell proliferation through glutathione depletion^20,25^, induce cytoprotective responses including hemeoxygenase-1^21^, and/or promote regulatory T cell phenotypes^17^. Notably, genetic reduction of HAAO extends lifespan in *C. elegans* and mice through mechanisms attributed at least in part to elevated 3HAA^24^. In the context of RENCA tumors, these prior observations motivate testable hypotheses linking HAAO disruption to altered redox and/or immune interactions, but the specific mediators remain unresolved in the current dataset.

In contrast to *Haao, Kmo* and *Qprt* knockouts produced more consistent phenotypes across experimental contexts. Both knockouts reduced migration in transwell assays and decreased colony size *in vitro*, and generated fewer metastatic events and increased survival in RCC mouse models relative to wild type cells. *Qprt* cells also resulted in lower tumor burden, while *Kmo* cells trended toward higher tumor burden (significant at 10 days post injection). The consistency between *in vitro* and *in vivo* results suggests that intrinsic tumor cell changes are sufficient to impair progression, although we cannot exclude contributions from immune or stromal components. Based on pathway topology, both *Kmo* and *Qprt* knockouts are predicted to reduce flux through the NAD^+^ biosynthesis branch: KMO initiates the hydroxylation sequence leading toward QA, while QPRT catalyzes the final step in de novo NAD^+^ production. We did not directly measure NAD^+^ levels in these experiments, but the convergent phenotypes are consistent with impaired NAD^+^ synthesis as a shared mechanism. NAD^+^ supports glycolysis, mitochondrial function, DNA repair through PARP enzymes, and epigenetic regulation through sirtuins—processes critical for cancer cell proliferation and stress resistance^15,16^. If NAD^+^ availability is indeed reduced in *Kmo* and *Qprt* knockout cells, this could explain their impaired proliferative and migratory capacity.

The convergent effects of *Kmo* and *Qprt* knockouts may also result in accumulation of upstream metabolites with distinct biological activities: increased KYN in the case of *Kmo*, and increased QA in the case of *Qprt*. KYN is an AhR ligand with documented immunosuppressive properties^11,12^, while QA is an NMDA receptor agonist that can induce cellular stress^22,23^, effects considered tumor-promoting in other contexts. One possibility is that NAD^+^ depletion is the dominant effect, overriding any pro-tumorigenic contributions from metabolite accumulation. Alternatively, accumulated metabolites may not reach concentrations sufficient to influence tumor behavior, or their effects may require conditions not present in our experimental systems. Metabolite profiling of *Kmo* and *Qprt* knockout cells and tumors would be needed to distinguish among these possibilities.

These findings have potential implications for therapeutic targeting of the kynurenine pathway in RCC. Clinical efforts to block the KP have focused predominantly on IDO1 inhibitors, with the rationale that restoring tryptophan availability would relieve immunosuppression in the tumor microenvironment. However, IDO1 inhibitors have shown limited efficacy in clinical trials, including the phase III failure of epacadostat in combination with pembrolizumab^38^. Our data suggest that targeting downstream enzymes in the KP could provide anti-tumor effects that differ from, and potentially complement, IDO1 inhibition, though whether this reflects differences in metabolite accumulation, NAD^+^ availability, or other mechanisms remains to be determined. The observation that all three knockouts extended survival, despite producing divergent effects on *in vitro* aggressiveness, indicates that multiple intervention points KP can impair RCC progression.

Several limitations should be considered when interpreting these results. First, we used a single RCC model (RENCA) in a syngeneic mouse system; whether these findings generalize to other RCC subtypes or to human disease is unknown. Second, we did not directly measure NAD^+^ or kynurenine pathway metabolites, so our mechanistic interpretations are based on predicted pathway topology rather than confirmed metabolic changes. Third, we did not characterize immune infiltration or activation state in knockout tumors, leaving open the question of whether immune-mediated mechanisms contribute to the reduced tumor burden observed with *Haao* knockout. Finally, we used complete genetic ablation rather than pharmacological inhibition; small-molecule inhibitors may produce different effects if they incompletely block enzyme activity or do not affect non-enzymatic protein functions.

Several follow-up directions would help resolve mechanisms suggested by the current findings. Direct metabolite profiling of each knockout cell lines and resulting tumors could confirm whether the shifts in 3HAA, KYN, QA, and/or NAD^+^ accompany each gene knockout. Characterization of tumor-microenvironment features (e.g., immune cells, stress response) could help explain discordant in vitro versus *in vivo* behavior observed with Haao knockout. Rescue experiments—for example, supplementing HAAO knockout cells with QA or NAD^+^ precursors—could help distinguish upstream vs. downstream metabolites as mediators of the observed phenotypes. determine whether restoring downstream metabolites reverses the observed phenotypes. These experiments are beyond the scope of the present study but represent clear next steps for understanding how KP nodes influence tumor progression.

## Conclusion

Our findings reveal enzyme- and context-dependent effects of KP disruption in RCC. *Haao* knockout increased migratory capacity and colony size *in vitro* but reduced tumor burden and metastasis *in vivo*, while *Kmo* and *Qprt* knockouts consistently impaired tumor-promoting phenotypes across both contexts. All three knockouts extended survival in a syngeneic mouse model, indicating that the NAD^+^ biosynthesis branch of the kynurenine pathway contributes to RCC progression. The divergent behavior of *Haao* knockout cells highlights that *in vitro* aggressiveness does not reliably predict *in vivo* tumorigenicity, particularly when metabolic pathways with pleiotropic effects are perturbed. These results suggest that individual KP enzymes represent distinct therapeutic targets with different biological consequences, and that the development of enzyme-specific inhibitors warrants further investigation in RCC and other cancers where kynurenine metabolism is dysregulated.

## Notes

### Competing Interest Statement

The authors have declared no competing interest.

### Summary of Updates

Minor edits and corrections in preparation for journal submission.

### Funding

- United States Department of Defense: DOD W81XWH2110540

