## Supplemental data for "Kynurenine Metabolism Mediates Tumor Progression in Renal Cell Carcinoma"

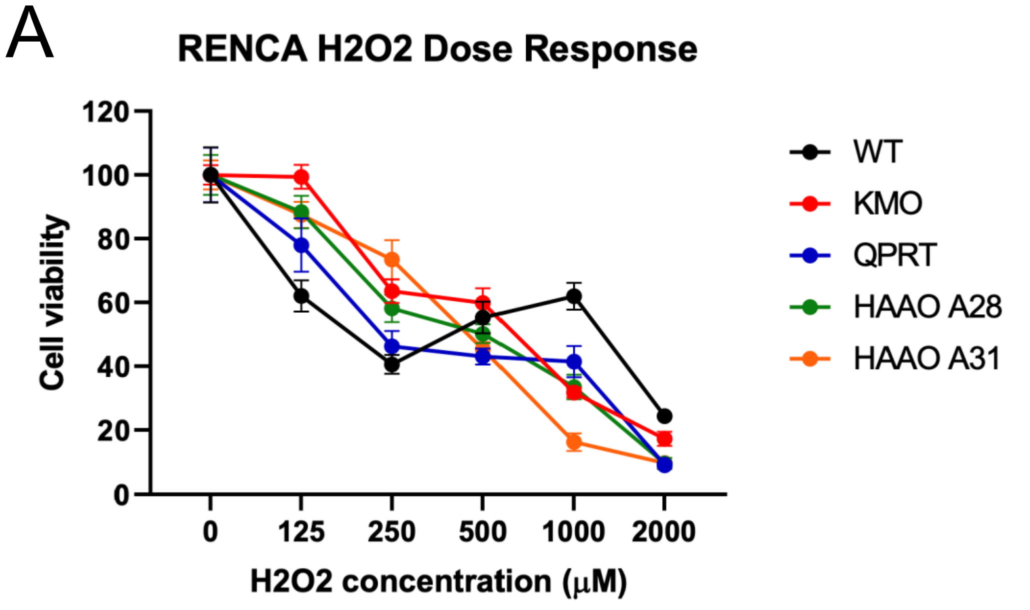


**Supplemental figure S1:** There is no significant difference between cell lines in response to hydrogen peroxide ($H_2O_2$) dose response.

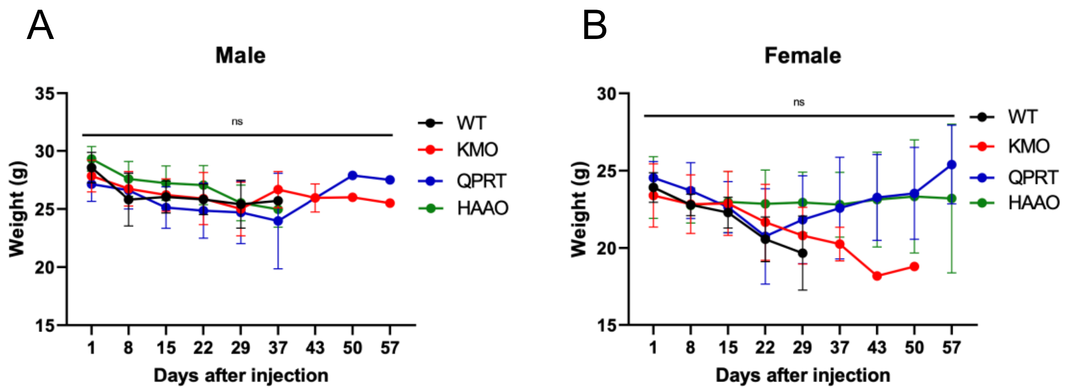


**Supplemental figure S2:** There was no significant change in weight in male (A) or female (B) mice during treatment.

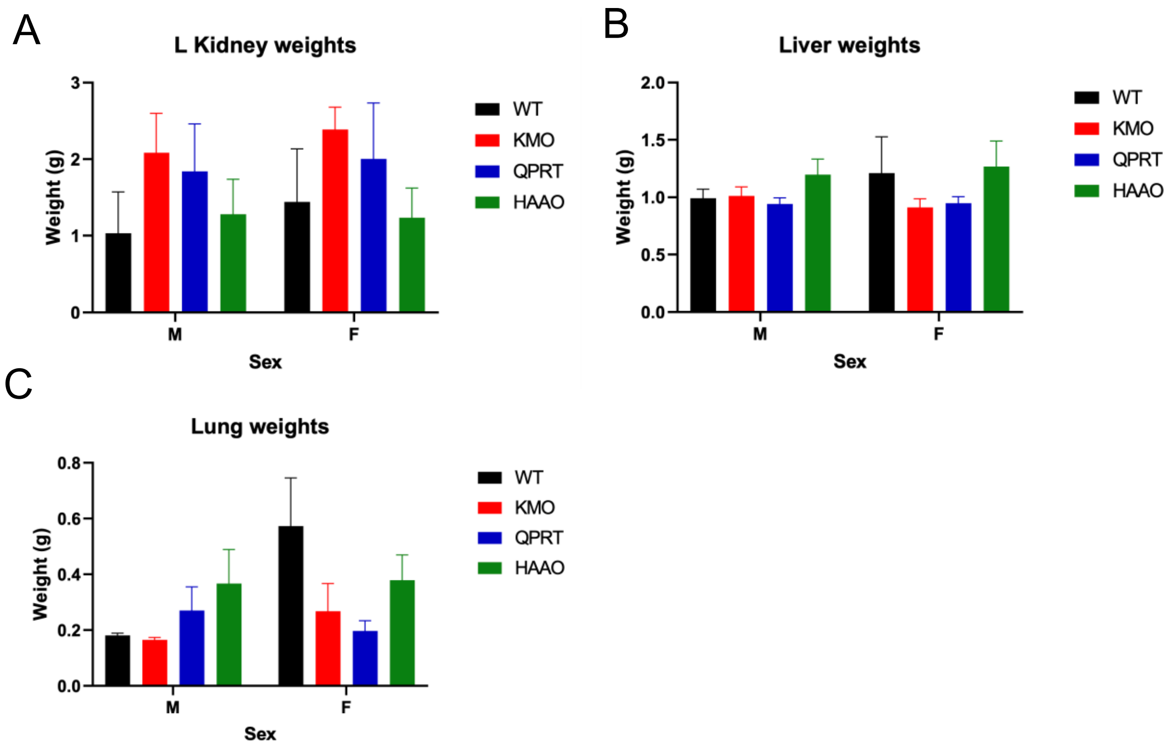


**Supplemental figure S3:** There was no significant difference between groups in left kidney weight (A), liver weight (B), or lung weights (C).

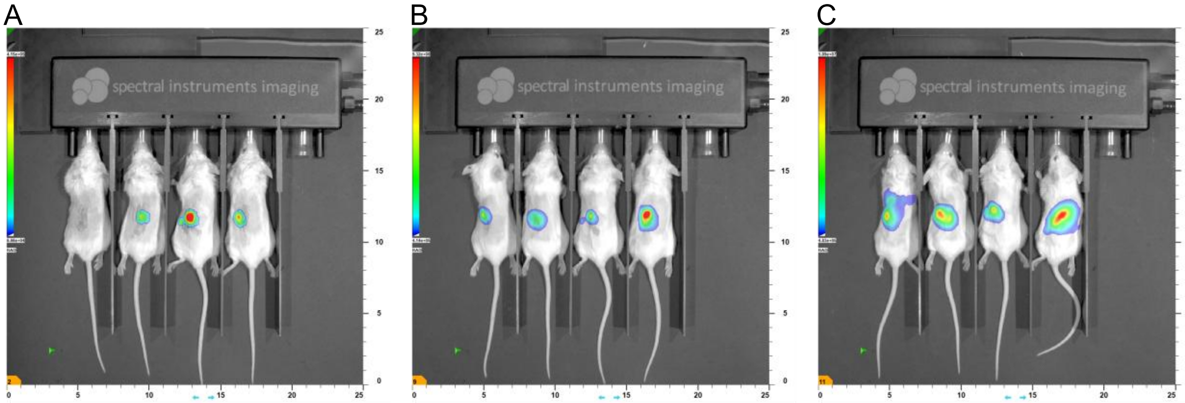



**Supplemental figure S4:** Representative images of in vivo bioluminescence at day 3 (A), day 10 (B), and day 17 (C).

| | |
|---|---|
| Median survival | |
| WT | 30.50 |
| HAAO | 38.00 |
| Log-rank (Mantel-Cox) test | |
| Chi square | 29.03 |
| df | 1 |
| P value | <0.0001 |
| P value summary | **** |

| | |
|---|---|
| Median survival | |
| KMO | 38.00 |
| WT | 30.50 |
| Log-rank (Mantel-Cox) test | |
| Chi square | 15.04 |
| df | 1 |
| P value | 0.0001 |
| P value summary | *** |

| | |
|---|---|
| Median survival | |
| QPRT | 35.50 |
| WT | 30.50 |
| Log-rank (Mantel-Cox) test | |
| Chi square | 5.327 |
| df | 1 |
| P value | 0.0210 |
| P value summary | * |

**Supplemental figure S5:** Statistics of Kaplan-Meier survival curve.
